# Expansion and Environmental Drivers of Invasive Freshwater Molluscs in Catalonia

**DOI:** 10.64898/2026.01.05.697568

**Authors:** Rafel Rocaspana, Quim Pou-Rovira, Enric Aparicio

## Abstract

Freshwater ecosystems in the Mediterranean region are highly vulnerable to biological invasions, yet invasive macroinvertebrates remain less studied than other taxonomic groups. In Catalonia (NE Spain), non-native freshwater molluscs are increasingly widespread and may substantially alter ecosystem functioning. This study analyses the spatiotemporal expansion and environmental drivers of four invasive freshwater molluscs (*Physella acuta, Potamopyrgus antipodarum, Corbicula fluminea and Dreissena polymorpha*) across the Catalan River Basin District. Species occurrence data were compiled from regional and global biodiversity databases and long-term bio-monitoring programmes, harmonised and aggregated into a 10 × 10 km grid. Temporal trends in cumulative occupancy were used to quantify expansion rates, while Generalized Linear Models were applied to identify key environmental and anthropogenic correlates of species occurrence. Gastropods (*P. acuta* and *P. antipodarum*) exhibited rapid and extensive range expansion since the mid-2000s, whereas bivalves showed limited or negligible spread. Expansion patterns were partly influenced by increased sampling effort following the implementation of the EU Water Framework Directive. Species distributions were consistently associated with low-altitude, low-slope river reaches subject to high anthropogenic pressure. *Physella acuta* was strongly linked to degraded lowland habitats, *C. fluminea* to large, hydrologically stable river sections, and *P. antipodarum* was primarily constrained by channel slope. These results reveal a longitudinal zonation of invasion risk in Mediterranean rivers and highlight the combined role of hydromorphology and human pressure in shaping invasion dynamics. Maintaining long-term monitoring and protecting headwater and mid-mountain reaches are essential to limit further spread and safeguard native freshwater biodiversity.

## 1. Introduction

Inland aquatic ecosystems are among the most vulnerable biomes globally, currently facing a severe biodiversity crisis, with extinction rates that exceed trends recorded in terrestrial and marine environments (Strayer & Dudgeon, 2010; Reid *et al*., 2019). This decline is intensified in the Mediterranean region, where aquatic biota must contend with the synergistic effects of hydrological extremes—exacerbated by climate change—and chronic anthropogenic stress (Fathy, 2024). Within this context, biological invasions have emerged as a primary driver of biodiversity loss and ecosystem homogenization. The introduction of Invasive Alien Species (IAS) alters community structure and disrupts ecosystem functioning, a pattern evident across Europe (Vilà *et al*., 2009). The Iberian Peninsula, particularly regions such as Catalonia in northeastern Spain, represents a continental hotspot for freshwater invasions, with over 30 non-native fish species recorded and increasing colonization by invertebrates (Muñoz-Mas & García-Berthou, 2020; Soto *et al*., 2025).

Despite the growing recognition of biological invasions as a global threat, research efforts exhibit a distinct taxonomic bias. In the Iberian Peninsula, as in much of Europe, the impacts and distribution patterns of alien fish have been extensively studied, driven primarily by their relevance to recreational fisheries (Cobo *et al*., 2010). Conversely, aquatic macroinvertebrates have received comparatively less attention. Among alien invertebrates, freshwater molluscs constitute a major component of aquatic invasive fauna, accounting for approximately 16% of non-indigenous animal species in the Iberian Peninsula, with gastropods and bivalves being the most prevalent classes (Zamora-Marín *et al*., 2023).

Non-native bivalves and gastropods are a serious threat to freshwater integrity due to their high fecundity, rapid growth, tolerance to degraded water quality, and capacity to alter benthic food webs (Sousa *et al*., 2008; Ricciardi, A, 2015). Notable species include the Asian clam *Corbicula fluminea*, the New Zealand mudsnail *Potamopyrgus antipodarum*, the ramshorn snail *Physella acuta*, and the zebra mussel *Dreissena polymorpha*, frequently form co-occurring assemblages that thrive in degraded conditions. Although research has historically prioritized economically damaging bivalves like *Dreissena polymorpha* (Dölle & Kurzmann, 2020), gastropods such as *P. acuta* and *P. antipodarum* are widely established across the region, often associated with ecological impacts that remain poorly quantified despite being yet equally pervasive (Vinarski, 2017; Alonso, 2019). Catalonia (NE Spain) has historically acted as a primary gateway for these introductions; notably, the first Iberian record of *P. antipodarum* was recorded in the Llobregat Delta (Barcelona) in 1924 (Alonso, 2019). Concurrently, intense human activities—including agricultural runoff, urban wastewater discharge, and dam construction—create synergistic pressure that may favour invasion success (Varga *et al*., 2019; Peñuelas *et al*., 2021). Despite this, comprehensive assessments of molluscan invasions across this region, particularly their relationship with environmental and anthropogenic factors, remain limited.

This study aims to: (1) map and analyse the spatiotemporal distribution changes of exotic bivalves and gastropods in Catalan rivers between 2010 and 2024; and (2) identify environmental and anthropogenic correlates of their distribution using multivariate statistical models. The findings will contribute to improve predictive models of invasion risk in similarly stressed Mediterranean basins.

## 2. Materials and Methods

### 2.2. Study Area

The study was conducted in the Catalan River Basin District (CRBD), which covers approximately 16,500 km^2^ in northeastern Spain. The area comprises several small to medium-sized river basins and is managed by the Catalan Water Agency (Agència Catalana de l’Aigua, ACA). The region hosts over 300 freshwater monitoring stations operated by the ACA, providing long-term ecological and physicochemical data. Rivers in the region are predominantly Mediterranean, characterized by flashy hydrology, summer droughts, and winter floods. This study focused on major basins, including the Ter, Llobregat, Fluvià, and Muga (Fig. 1), based on data availability and the overall significance of these basins.

**Figure 1.**
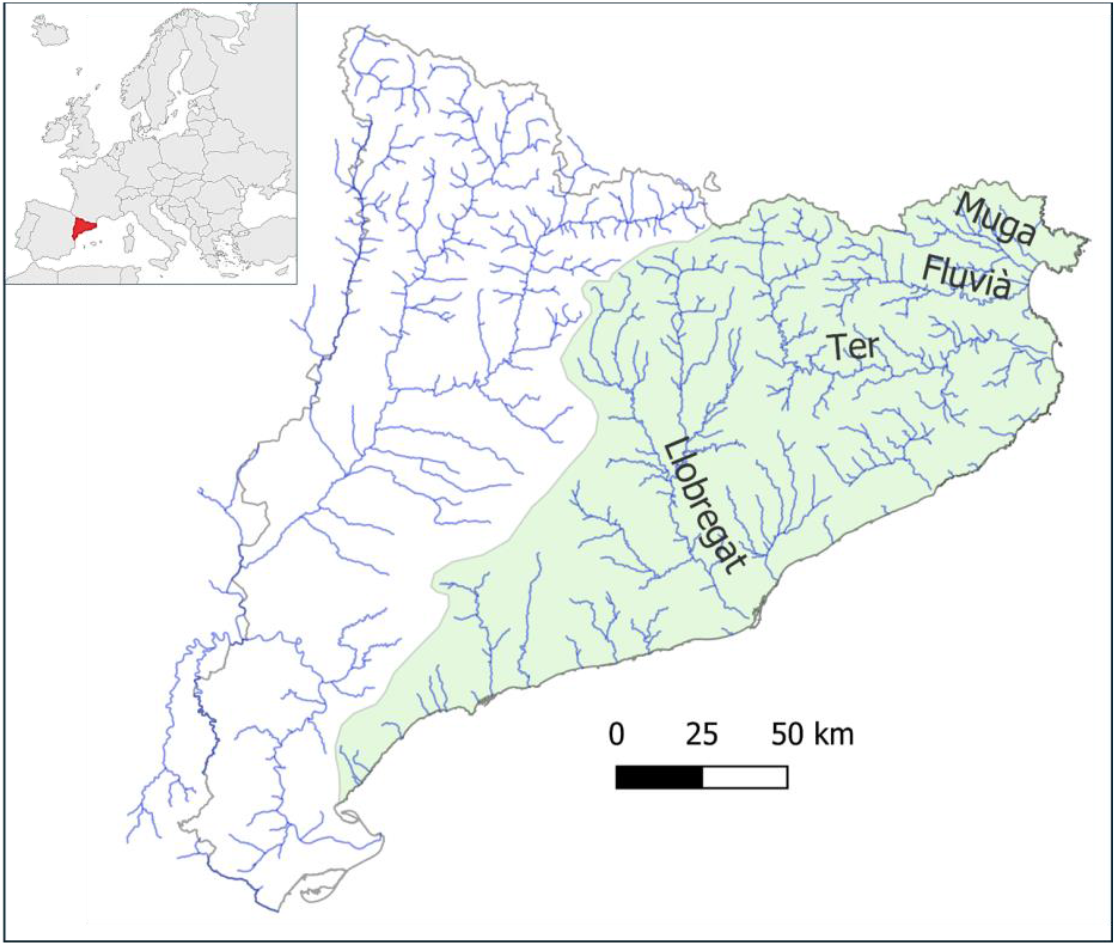
Geographic location of the study area within Europe. Catalonia is highlighted in red, and the main river basins within the Catalan River Basin District (background green layer) are indicated.

Catalonia’s rivers are highly modified: >60% are classified as having moderate to poor ecological status under the EU Water Framework Directive (ACA, 2023). Urban areas (e.g., Barcelona, Girona) and intensive agriculture (irrigated crops and livestock) contribute significant nutrient and contaminant loads. The area has experienced a mean warming of +1.3°C since 1980 (Lana *et al*., 2022), enhancing thermal suitability for warm-adapted exotic species.

### 2.2. Data Collection

Occurrence data for invasive freshwater bivalves and gastropods were compiled from multiple open-access biodiversity and environmental data repositories covering both global and regional scales. Records were obtained from the Global Biodiversity Information Facility (GBIF; https://www.gbif.org/), the EXOCAT database of exotic species of Catalonia (https://exocat.creaf.cat/), the Biodiversity Data Bank of Catalonia (BDBC; http://biodiver.bio.ub.es/bdbc/), and the biological monitoring datasets of the ACA. Data extraction from GBIF was performed by querying for the target taxonomic groups within the geographic extent of Catalonia, including all available records with georeferenced information and valid taxonomic identification. Regional occurrence records were complemented with data from EXOCAT and the BDBC to improve coverage of locally recorded invasive species and to include records not indexed in global repositories. Additional occurrences and ecological information were obtained from the ACA bio-monitoring programme, which provides standardized biological and environmental data collected under official river basin monitoring protocols.

The species for which data were collected were *C. fluminea, P. antipodarum, P. acuta*, and *D. polymorpha*. Other introduced mollusc species are present, but either have very restricted or localized distributions (such as *Planorbella duryi* and *Sinanodonta woodiana*) or show very limited occurrence in rivers (e.g., *Pomacea* spp.). Consequently, these species were excluded from the study. All retrieved records were integrated into a unified database and harmonised using standard Darwin Core terms to ensure consistency across sources. Records lacking geographic coordinates or with clearly erroneous spatial information were excluded from further analyses. The final dataset therefore represents a harmonised compilation of invasive freshwater bivalve and gastropod occurrences in Catalonia, suitable for subsequent analyses of spatial distribution, temporal trends in range dynamics, and relationships with environmental variables and anthropogenic pressures.

### 2.3. Spatial Analysis

To perform a standardised spatial analysis, geographic coordinates were projected using QGIS (QGIS Development Team, 2025) into the Universal Transverse Mercator (UTM) coordinate system, specifically Zone 31N, which is the standard projection for the study area in Catalonia. A grid-based approach was adopted to mitigate sampling bias and account for spatial uncertainty. Occurrences were aggregated into a 10×10 km UTM grid. This resolution is consistent with international standards for calculating the Area of Occupancy (AOO) in biodiversity assessments.

To evaluate the expansion or contraction of the species’ ranges, we calculated the cumulative occupancy. Unlike annual occupancy, which can be heavily influenced by fluctuations in sampling effort, cumulative occupancy represents the total number of unique 10 x 10 km grid cells where a species has been recorded up to a given year. This metric is particularly robust for monitoring the colonisation process of invasive or expanding species.

The rate of expansion was quantified using linear regression models, where the year was the independent variable and the cumulative number of occupied grid cells was the dependent variable. The slope of the regression line was used as a proxy for the expansion rate (new grid cells colonised per year). Statistical significance was determined using a *p*-value threshold of α = 0.05. A significant positive slope indicated an increasing trend, a slope close to zero denoted stability.

### 2.4. Environmental and anthropogenic correlates

To investigate the association between environmental and anthropogenic factors and the occurrence of exotic molluscs, we compiled several site-specific hydrological and climatic descriptors. Catchment attributes—including altitude (m a.s.l.), drainage area (km^2^), and reach slope (m km^−1^)—were extracted from publicly available geographic datasets, and mean air temperature was obtained from the BioClim database (Karger *et al*., 2017). To assess environmental degradation at each site, we used an Environmental Disturbance Score (EDS) that integrates multiple categories of anthropogenic pressure: nutrient enrichment, non-natural land use, channel morphology alteration, connectivity disruption, and flow regulation. Disturbance levels for each category were assigned based on expert judgement and adapted from the classification framework proposed by Faro *et al*. (2024). Each variable was scored according to its deviation from minimally disturbed conditions, ranging from 1 (no deviation) to 4 (highly degraded). For each site, the five individual scores were summed to produce a composite EDS.

Differences in environmental and degradation variables between sites of presence and absence were initially assessed using the non-parametric Mann-Whitney *U* test, as most variables did not follow a normal distribution.

To identify the main drivers of species distribution, Generalized Linear Models (GLM) with a binomial error distribution and a logit link function were constructed. Prior to modelling, multicollinearity among predictors was evaluated using the Variance Inflation Factor (VIF). Variables exhibiting high redundancy were removed, retaining a final subset of predictors with VIF < 2 to ensure model stability. All predictor variables were standardized (mean = 0, SD = 1) before analysis to allow for a direct comparison of effect sizes through the estimated regression coefficients. The significance of predictors was assessed using the Wald test (*p* < 0.05), and the overall model fit was evaluated using McFadden’s Pseudo-*R*^2^. To visualize the independent effect of the most significant predictors, we calculated marginal response curves based on the fitted GLMs. Predicted probabilities were estimated across the range of the focal variable while holding all other predictors constant at their mean values. All statistical analyses were conducted in R version 4.4.0 (R Core Team, 2025).

## 3. Results

The cumulative spatial expansion of aquatic molluscs monitored between 1956 and 2025 reveals different colonisation velocities, with a clear divergence between rapidly spreading gastropods and relatively static bivalve species (Fig. 2).

**Figure 2.**
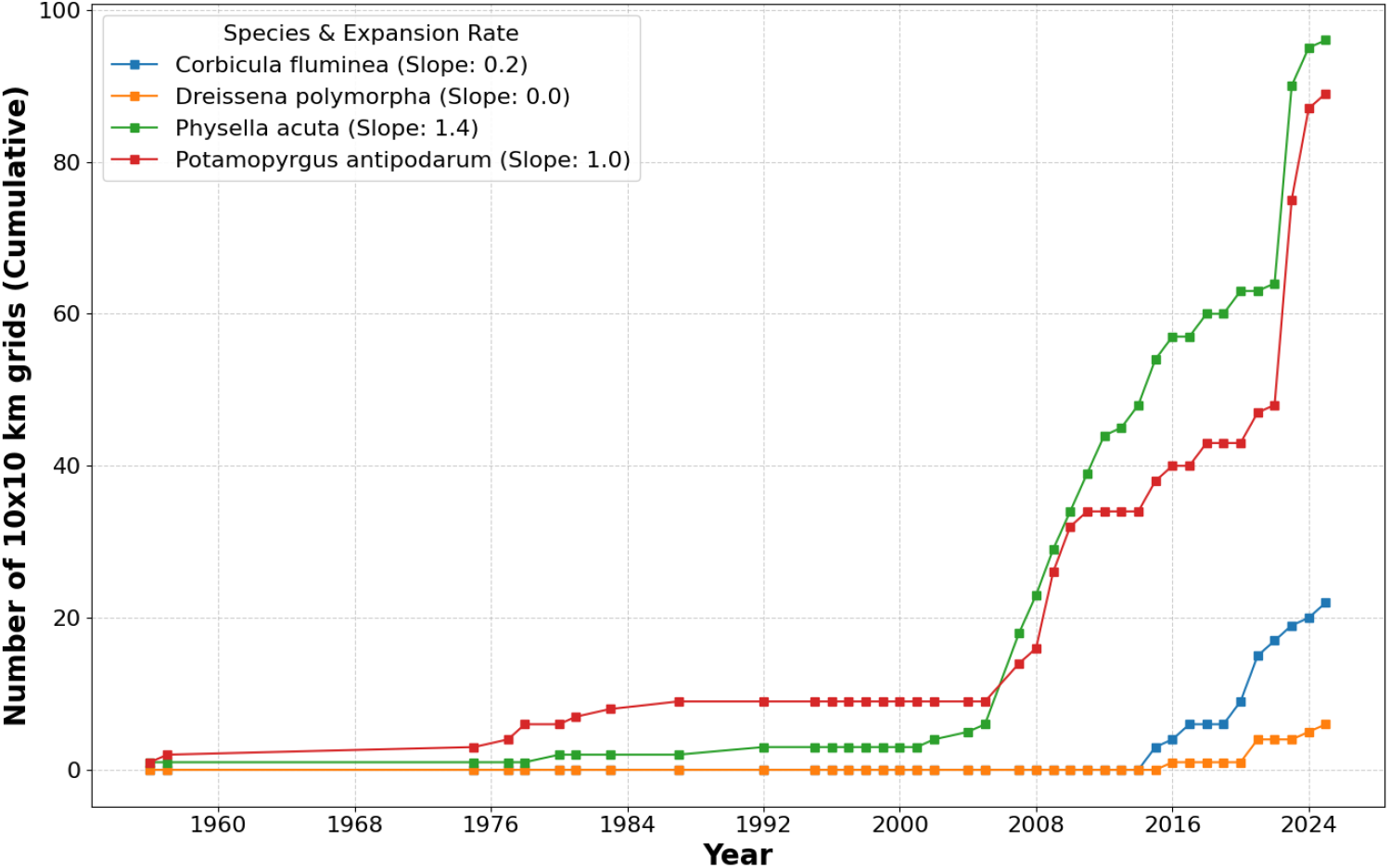
Temporal expansion trends of the invasive species *Corbicula fluminea, Dreissena polymorpha, Physella acuta* and *Potamopyrgus antipodarum* in the Catalan River Basin District. The plot displays the cumulative increase in the Area of Occupancy (AOO), measured as the number of occupied 10×10 km UTM grid cells over time. The slope of the linear regression is provided in the legend.

Gastropods exhibited the most pronounced expansion dynamics. *Physella acuta* showed the fastest spread (slope = 1.4); after a prolonged lag phase, its spatial occupancy increased sharply around 2006–2008. A similar pattern was observed for *P. antipodarum* (slope = 1.0), which remained restricted until the mid-2000s before entering a phase of rapid expansion. By the end of the study period (2025), these two taxa reached the highest occupancy values, exceeding 90 and 80 grids, respectively (Fig.3).

In contrast, bivalve species displayed much lower expansion velocities. *Corbicula fluminea* showed a late-onset expansion, beginning approximately in 2015, resulting in a moderate progression slope of 0.2 and a final occupancy of 22 grids (Fig. 3). Meanwhile, *D. polymorpha* exhibited negligible territorial growth across the entire time series (slope: 0.0), remaining restricted to fewer than 10 grids by 2025 (Fig. 3).

**Figure 3.**
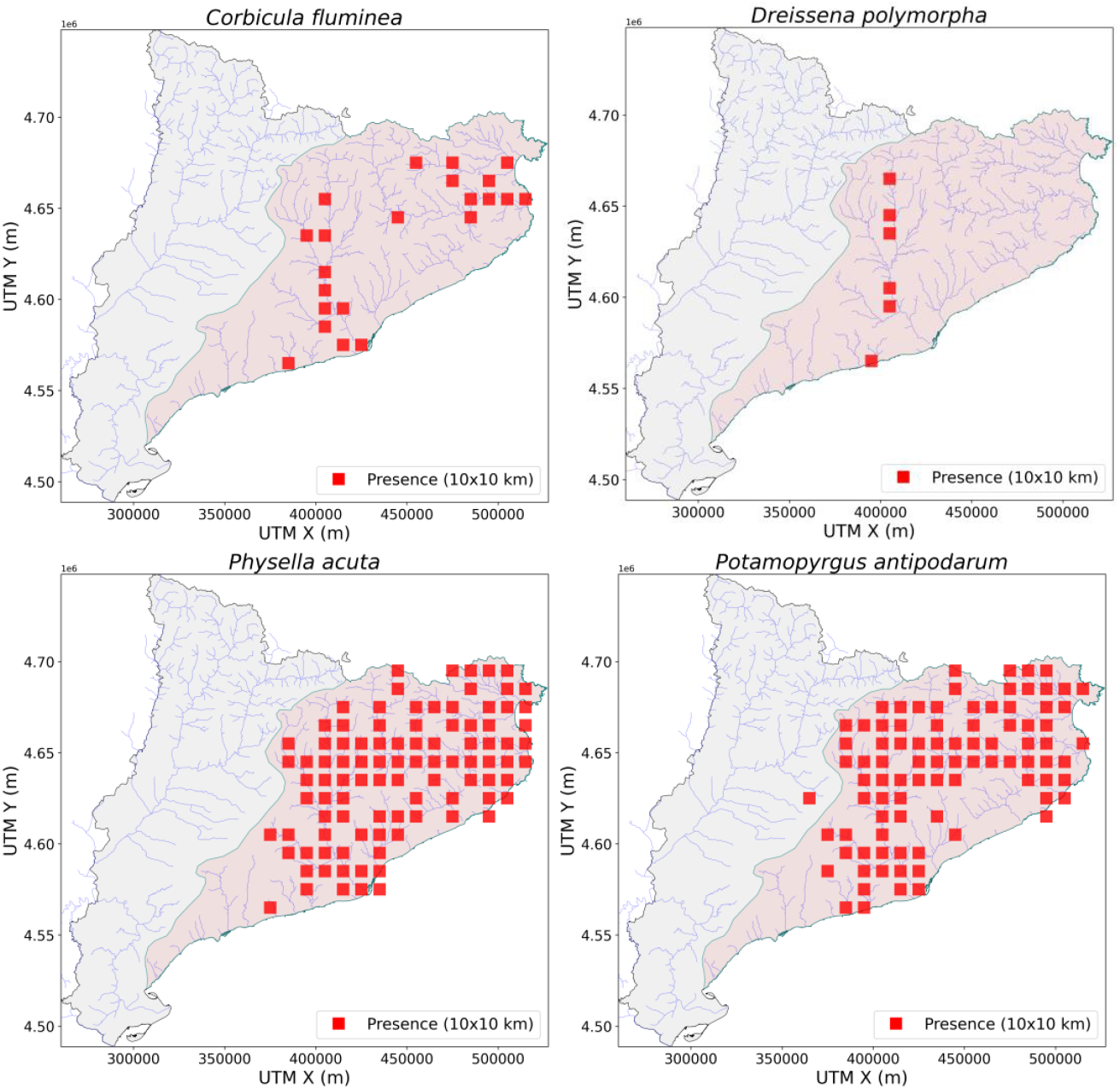
Distribution maps of the invasive species *Corbicula fluminea, Dreissena polymorpha, Physella acuta* and *Potamopyrgus antipodarum* in the Catalan River Basin District (background red layer). Presence is recorded in 10×10 km UTM grid cells (Zone 31N), integrating all available historical and current records.

Univariate analyses showed that the three most widespread species (excluding *D. polymorpha* due to its low sample size) were consistently associated with distinct environmental and hydromorphological conditions compared with sites where they were absent (Table 1). Overall, *P. antipodarum, C. fluminea*, and *P. acuta* tended to occur in river reaches at lower altitudes, with gentler slopes, and under higher levels of anthropogenic pressure (EDS index). Among them, *C. fluminea* displayed the clearest habitat segregation, being largely confined to sites with much larger catchment areas and located farther from the river source than the other two species. Although *P. acuta* and *P. antipodarum* occupied broadly similar environmental ranges, *P. acuta* showed a stronger association with warmer mean water temperatures and higher levels of anthropogenic disturbance.

**Table 1.** Summary statistics (mean ± SD) of environmental variables and anthropogenic pressures (EDS) for presence and absence sites of *Corbicula fluminea, Physella acuta* and *Potamopyrgus antipodarum*, including *Mann-Whitney U* test results.

| Variable | Species | Presence (Mean ± SD) | Absence (Mean ± SD) | p-value |
| --- | --- | --- | --- | --- |
| EDS | *P. antipodarum* | 12.70 ± 3.90 | 9.95 ± 2.70 | 0.003 |
| | *C. fluminea* | 17.05 ± 3.52 | 11.69 ± 3.45 | <0.001 |
| | *P. acuta* | 13.05 ± 3.79 | 9.71 ± 2.99 | <0.001 |
| Altitude (m) | *P. antipodarum* | 324.80 ± 267.11 | 532.89 ± 410.45 | 0.024 |
| | *C. fluminea* | 134.00 ± 161.07 | 379.51 ± 295.84 | <0.001 |
| | *P. acuta* | 282.81 ± 233.12 | 599.15 ± 360.98 | <0.001 |
| Catchment area (km<sup>2</sup>) | *P. antipodarum* | 374.00 ± 852.69 | 109.45 ± 317.79 | 0.001 |
| | *C. fluminea* | 1731.35 ± 1618.19 | 147.43 ± 297.80 | <0.001 |
| | *P. acuta* | 412.30 ± 899.33 | 81.98 ± 133.61 | 0.024 |
| Slope (%) | *P. antipodarum* | 14.03 ± 14.24 | 30.78 ± 23.71 | <0.001 |
| | *C. fluminea* | 6.52 ± 8.66 | 17.33 ± 16.87 | <0.001 |
| | *P. acuta* | 12.92 ± 13.70 | 26.83 ± 21.03 | <0.001 |
| Mean temperature (°C) | *P. antipodarum* | 12.72 ± 2.71 | 11.87 ± 2.46 | 0.073 |
| | *C. fluminea* | 14.25 ± 1.25 | 12.39 ± 2.76 | <0.001 |
| | *P. acuta* | 12.94 ± 2.74 | 11.42 ± 2.12 | <0.001 |

Generalized Linear Models (GLMs) identified distinct significant predictors for the distribution of each species after controlling for covariate interdependencies (Table 2; Fig 4). For *P. acuta* (Pseudo-*R*^*2*^ = 0.28), occurrence was mainly explained by a strong positive association with anthropogenic pressure (EDS; *p* < 0.01) and a significant negative relationship with altitude (*p* < 0.01). In contrast, the distribution of *C. fluminea* (Pseudo-*R*^2^ = 0.46) was predominantly explained by hydromorphological scale, with catchment area being the only significant positive predictor (*p* < 0.01) in the multivariate model, while the effect of EDS was marginally significant. Finally, the model for *P. antipodarum* (Pseudo-*R*2 = 0.16) identified channel slope as the primary limiting factor, showing a significant negative association (*p* < 0.05), suggesting that hydraulic conditions, rather than altitude or anthropogenic pressure alone, significantly constrain its local distribution.

**Table 2.** Results of the Generalized Linear Model (GLM, binomial distribution) explaining the occurrence probability of *Corbicula fluminea, Physella acuta* and *Potamopyrgus antipodarum*. Predictor variables were standardized (z-score) prior to analysis to allow direct comparison of effect sizes.

| Predictor | Species | $\beta$ | SE | Z-value | p-value | Odds Ratio | Model Fit (Pseudo-R<sup>2</sup>) |
| --- | --- | --- | --- | --- | --- | --- | --- |
| EDS | *P. antipodarum* | 0.724 | 0.4 | 1.803 | 0.071 | 2.06 | 0.162 |
|  | *C. fluminea* | 0.765 | 0.42 | 1.834 | 0.067 | 2.15 | 0.455 |
|  | *P. acuta* | 1.029 | 0.36 | 2.892 | 0.004 | 2.80 | 0.275 |
| Altitude | *P. antipodarum* | -0.339 | 0.47 | -0.725 | 0.468 | 0.71 | 0.162 |
|  | *C. fluminea* | -0.733 | 0.64 | -1.144 | 0.253 | 0.48 | 0.455 |
|  | *P. acuta* | -0.945 | 0.34 | -2.800 | 0.005 | 0.39 | 0.275 |
| Catchment area | *P. antipodarum* | -0.132 | 0.59 | -0.225 | 0.822 | 0.88 | 0.162 |
|  | *C. fluminea* | 1.223 | 0.44 | 2.764 | 0.006 | 3.40 | 0.455 |
|  | *P. acuta* | -0.005 | 0.77 | -0.007 | 0.995 | 0.99 | 0.275 |
| Slope | *P. antipodarum* | -0.550 | 0.25 | -2.178 | 0.029 | 0.58 | 0.162 |
|  | *C. fluminea* | 0.07 | 0.53 | 0.133 | 0.895 | 1.07 | 0.455 |
|  | *P. acuta* | -0.180 | 0.24 | -0.762 | 0.446 | 0.84 | 0.275 |
| Mean temperature | *P. antipodarum* | -0.218 | 0.64 | -0.343 | 0.732 | 0.80 | 0.162 |
|  | *C. fluminea* | 0.25 | 0.62 | 0.405 | 0.685 | 1.28 | 0.455 |
|  | *P. acuta* | -0.005 | 0.38 | -0.015 | 0.988 | 0.99 | 0.275 |

**Figure 4.**
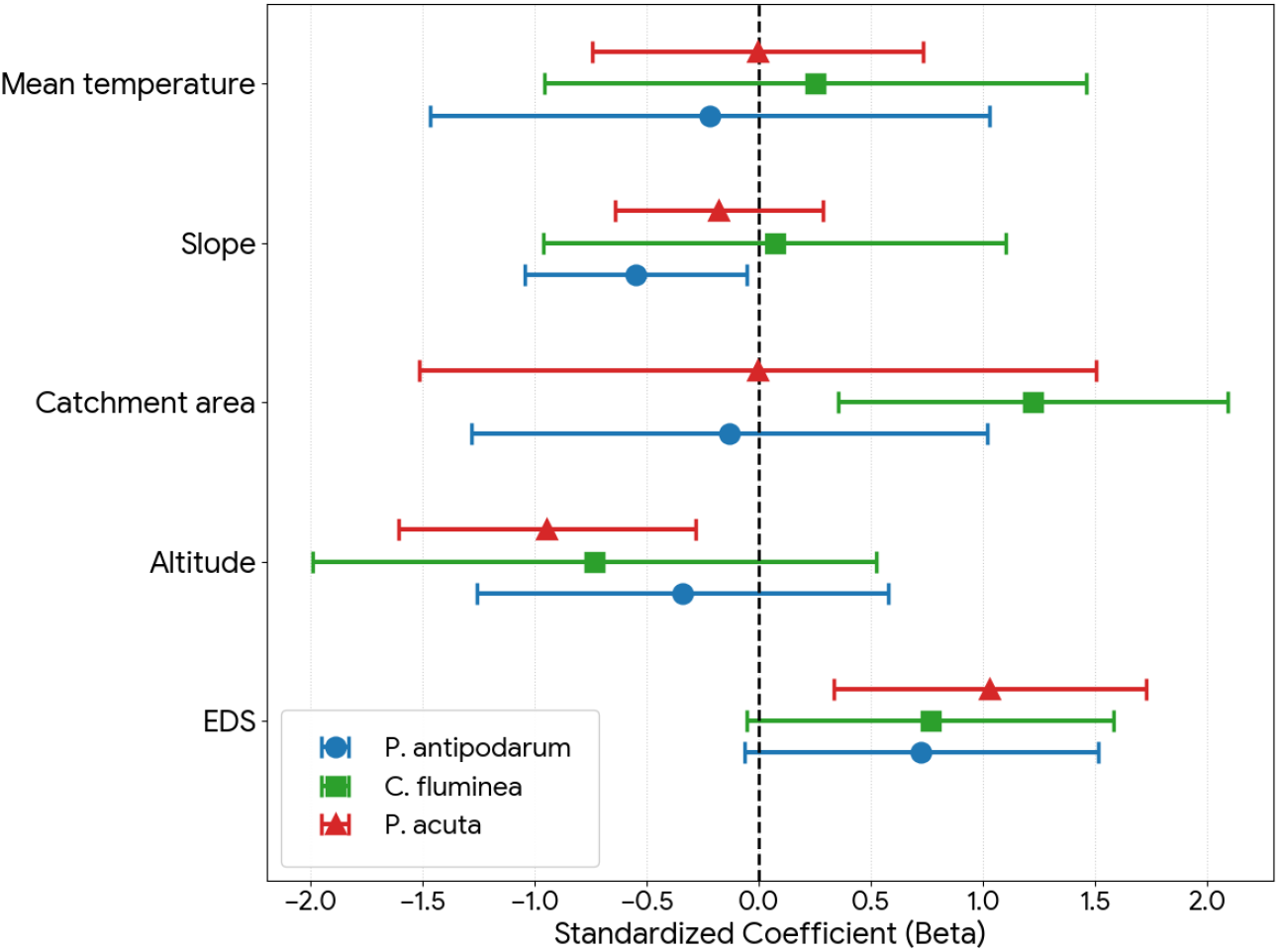
Forest plot summarizing the standardized regression coefficients (β) from Generalized Linear Models (GLMs) assessing the environmental drivers of *Potamopyrgus antipodarum, Corbicula fluminea*, and *Physella acuta* occurrence. Points represent the estimated mean effect size for each predictor, and horizontal error bars indicate the 95% confidence intervals. Predictors are considered statistically significant (*p* < 0.05) when their error bars do not intersect the vertical dashed line at zero. Positive coefficients indicate a positive association with species presence.

The analysis of marginal effects revealed distinct environmental constraints for each species (Fig. 5). *Potamopyrgus antipodarum* showed a sharp decline in occurrence probability as channel slope increased, suggesting a limitation by flow velocity in steep reaches. In contrast, *C. fluminea* exhibited a sigmoidal response to catchment area, being virtually absent in headwaters (< 100 km^2^) and peaking in large river sections. Finally, *P. acuta* displayed a strict altitudinal limit, with the probability of occurrence declining sharply as altitude increased.

**Figure 5.**
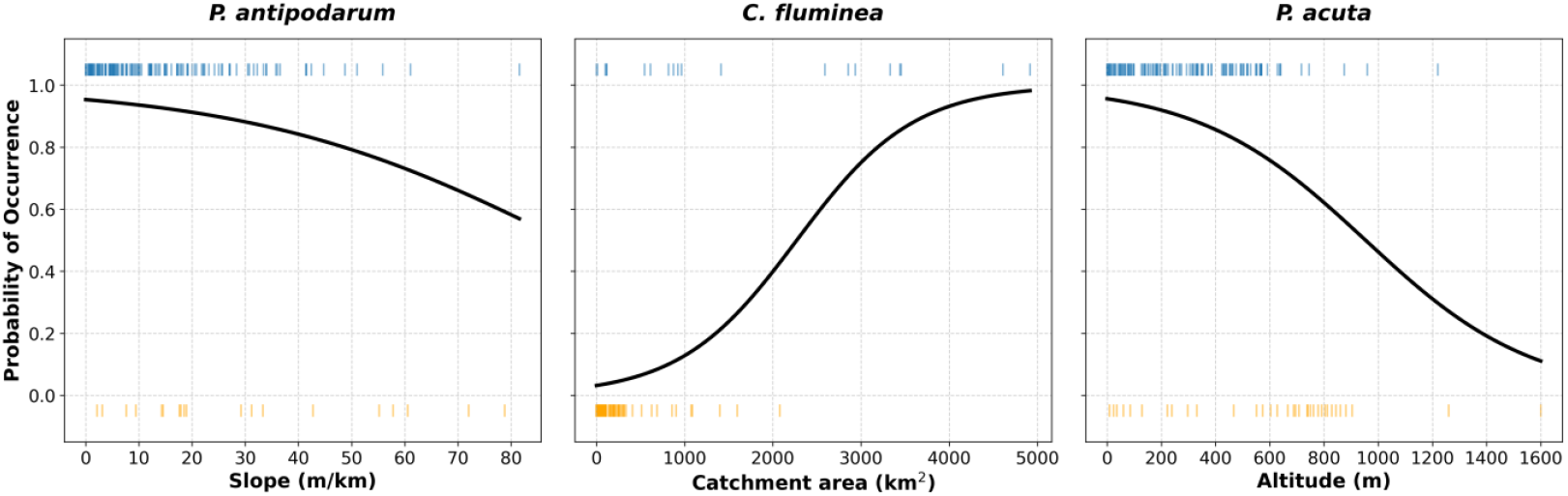
Response curves showing the predicted probability of occurrence for *Potamopyrgus antipodarum, Corbicula fluminea*, and *Physella acuta* along their most significant environmental gradients. The solid black lines represent the logistic regression models (GLM), estimating the marginal effect of each predictor while holding other variables constant at their mean values. The rug plots at the top (blue) and bottom (orange) of each panel indicate the observed presence and absence of the species, respectively.

## 4. Discussion

The spatiotemporal analysis of invasive freshwater molluscs in Catalonia highlights a clear contrast in colonisation dynamics between gastropods and bivalves. The results indicate that gastropods, specifically *P. acuta* and *P. antipodarum*, show the fastest and most expansive spread, ultimately occupying much of the Catalan River Basin District. In comparison, bivalve expansion is far more limited: *C. fluminea* shows a moderate but delayed increase in range, whereas *D. polymorpha* has exhibited almost no spatial growth, remaining confined to specific areas below reservoirs. However, interpreting these expansion curves requires careful consideration of sampling effort. The inflection points observed in our temporal data, particularly the sharp increase in species detection around 2006–2008, coincide with the systematic implementation of river monitoring programmes in Catalonia mandated by the EU Water Framework Directive (Munné *et al*., 2016). This suggests that a significant portion of the apparent sharp spread may be attributed to a sampling artefact rather than biological expansion alone. Before the WFD implementation, aquatic macroinvertebrates received comparatively less attention than other groups (e.g. fish) in the Iberian Peninsula. Consequently, the historical distribution of these species may be underestimated. Despite this sampling bias, the underlying biological trend remains evident. The rapid expansion of *P. acuta* and *P. antipodarum* in Catalonia aligns with broader patterns reported across the Iberian Peninsula and Central Europe, where invertebrates show no signs of saturation compared to fish species (Haubrock *et al*., 2023).

Our results demonstrate a convergent pattern among the studied invasive molluscs: a generalized preference for low-altitude reaches characterized by high anthropogenic pressure. The positive association between the EDS index and species occurrence, particularly for *P. acuta*, is consistent with the passenger model of invasion: habitat degradation weakens native communities and frees resources, thereby facilitating the establishment of non-native species (MacDougall & Turkington, 2005; Didham *et al*., 2007). *Physella acuta* emerged as a clear specialist of degraded, lowland habitats, with occurrence strongly negatively correlated with altitude and positively associated with anthropogenic pressure. Therefore, the species can serve as an indicator of environmental degradation. This pattern is consistent with a species adapted to nutrient-enriched and often hypoxic waters, where its ability to breathe atmospheric air likely confers a competitive advantage over less tolerant taxa (Koopman *et al*., 2016; Collado *et al*., 2025). Regarding *C. fluminea*, our Generalized Linear Models identified catchment area as the main predictor, effectively restricting the species to larger, more stable river sections. This finding aligns with previous studies showing that *C. fluminea* is favoured by stable water volumes and by sediments that remain well oxygenated. These conditions support benthic recruitment and the survival of pediveliger larvae, and are typically lacking in headwater reaches (Sousa *et al*., 2008; Modesto *et al*., 2025). In contrast to the other species, *P. antipodarum* was significantly constrained by channel slope, showing a preference for low-gradient reaches. Despite its high colonisation potential via parthenogenesis (Alonso & Castro-Díez, 2008), steep, high-velocity reaches constitute a primary limiting factor for its upstream expansion in high-gradient Mediterranean streams. This pattern is consistent with continental-scale analyses showing a decline in the share of non-native species with increasing elevation and steep channel gradients, which together limit the success of many freshwater invaders in headwater reaches (Haubrock *et al*., 2023). Therefore, the spatial segregation observed in this study suggests a longitudinal zonation of invasion risk and reinforces the view that headwater streams can act as refugia for native species by limiting the establishment and spread of non-native taxa (Boon *et al*., 2023). However, as climate change progresses, the thermal and hydrological barriers protecting these headwaters may weaken, potentially facilitating the upstream colonisation of thermophilic invaders (Khaliq *et al*., 2024).

In our study, *D. polymorpha* showed no evidence of expanding within the lotic waters of the CRBD, remaining confined to downstream reaches below reservoirs where established populations persist. This restricted distribution is consistent with patterns reported in other river systems, where upstream impoundments function as primary sources of larval dispersal, enabling downstream colonization but limiting broader spread due to hydrological and environmental constraints (Olson *et al*., 2018). These dynamics highlight the dependence of *D. polymorpha* populations in flowing waters on a continuous upstream supply of propagules and underscore the importance of reservoir management in mitigating invasive spread across river networks.

The relationship between sampling effort and species detection has direct management consequences. The apparent surge in species occurrence following the implementation of the WFD demonstrates that sporadic monitoring is insufficient for accurate invasion risk assessment. Because the detectability of aquatic invertebrates is heavily influenced by sampling intensity (Larras & Usseglio-Polatera, 2020), a cessation or reduction in monitoring frequency could cause a serious underestimation of future range expansions, delaying management responses and reducing the likelihood of successful containment or eradication. Therefore, it is imperative to maintain, and where possible increase, the intensity of monitoring programmes. Long-term time series are essential to distinguish between actual range expansions and artefacts of sampling effort. In addition, management should reflect the longitudinal zonation of invasion risk, prioritising biosecurity and early-detection efforts in lowland reaches with high human pressure to contain established populations, while strictly protecting the hydrological integrity of mid-mountain transition zones to maintain them as invasion barriers.

## Acknowledgments

The authors thank the Agència Catalana de l’Aigua for providing access to the biological and environmental monitoring data that partially supported this research.

